# Accelerating Evolutionary Hill Climbs in Parallel Turbidostats

**DOI:** 10.1101/217273

**Authors:** Christopher N. Takahashi, Luis Zaman, Eric Klavins

**Affiliations:** School of Computer Science and Engineering, University of Washington, Seattle; Center for the Study of Complex Systems, University of Michigan, Ann Arbor; Department of Electrical Engineering, University of Washington, Seattle

**Keywords:** Continuous culture, Turbidostat, Optimal evolution, Evolution model

## Abstract

Evolution has been used to address many engineering problems. Within the context of metabolic engineering and synthetic biology, directed evolution has natural applications. However, most research concerning optimizing microbial evolution has been focused on library generation and screening, while accelerating evolutionary hill climbs and been largely ignored. Here, we develop a model to explore how population structure can accelerate evolutionary hill climbs. We show that by adjusting the population size, environmental challenge, and meta-population dynamics that the rate of evolution can be accelerated in parallel turbidostats. Our analyses leads to two surprising results: small populations are favored over conventionally large microbial populations, and propagating modest fitness improvements is favored over propagating mutants with large beneficial mutations. When combined with rational design and other optimization techniques our theory can accelerate strain development for applications such as consolidated bioprocessing, and bioremidation systems.

## Introduction

Computational and mathematical models of evolution have helped uncovered fundamental patterns and processes occurring in nature (1–5). As our theoretical understanding of evolutionary dynamics improves, our ability to harness and direct microbial evolution towards desirable engineering goals increases. One such goal is to improve the production of economically relevant biosynthetic byproducts such as the antimaliarial drug precursor, artemisinic acid(6). Realizing that cheating is a common evolutionary solution when selecting cells via indirect markers (e.g., producing the marker without paying the cost of making the desired byproduct), (7) devised a method of alternating selection and counter-selection that effectively curtails *de novo* cheats and ultimately improved byproduct yield. In other words, understanding likely evolutionary outcomes allowed the authors to short circuit evolution with carefully engineered environments.

Here we investigate how multiplexed turbidostats (*i.e.*, bioreactors that allow controlled microbial population sizes) we have previously developed (8) can be used to speed up adaptation during directed evolutionary *hill climbs*. We develop a mathematical model of evolutionary dynamics within a turbidostat. We then consider a multiplexed metapopulation of turbidostats and ask whether decisions about how populations are propagated (e.g., isolated populations vs propagating the most fit population to the rest of the turbidostats at various intervals) can optimize the rate of adaptation. Our analyses led to two surprising results: small populations are favored over conventionally large microbial populations, and propagating modest fitness improvements is favored over propagating mutants with large beneficial mutations.

We combine steady state continuous culture dynamics (*e.g.*, in turbidostats and chemostats) with mutational events into a simple stochastic hybrid systems (SHS) (9) model. By considering a continuous culture environment much of the complexity associated with natural microbial populations much of the modeling complexity is removed. As a result, we are able to analytically compute the time to appearance of a more fit mutant and the time it takes that mutant to achieve majority in the population. Thus, we can query the model for optimal initial environments and population sizes for single and double step evolutionary hill climbs.

Our analysis demonstrates that uniformly sized steps are preferable to a large step followed by a small step due to the disproportionate amount of time required for small fitness improvements to sweep. This leads us to a counterintuitive control policy where large fitness jumps are rejected in favor of smaller but more uniform fitness increases. By applying this policy to a simulated metapopulation of bioreactors, we show that we can increase the rate of evolution. Finally, we examine the effects of population size and number of bioreactors in the metapopulation on the effectiveness of our control policy.

## Results

### Continuous Culture Dynamics

Here, we consider the case of continuous culture with microbes growing in a constant volume of well mixed liquid media. We begin with a standard ODE model for mixed populations in continuous culture (10)

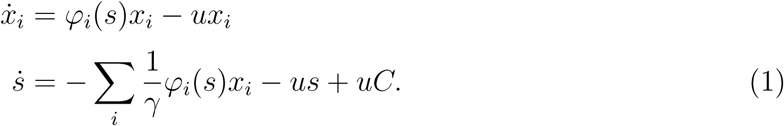

Here, *x*_*i*_ denotes the total amount of cells of genotype *i*. Individual cells divide at a rate *φ*_*i*_(*s*) for a limiting substrate concentration s. Cells and substrate are washed out with fresh media containing substrate at concentration *C* at a rate *u*. Finally, substrate is consumed at a rate of *γ* cells per unit of substrate.

### Turbidostat Evolutionary Dynamics

A turbidostat is a continuous culture device where the total cell population is held constant using feedback control(11). The resulting fixed population size allows us to make simplifying assumptions and enables us to apply previous work on fixed size populations by Moran (12). We begin by assuming the dilution rate *u*(*t*) is set by a feedback controller that ensures that the cell density is constant, which simplifies the resulting dynamics. While a complete model would include turbidostat feedback dynamics, such a system would only differ in the moments soon after inoculation as any reasonable controller will act on timescales much faster than the growth of microbes. We then extend the ODE model to a stochastic hybrid systems model to capture the random nature of mutations.

Mathematically, a fixed population size implies that

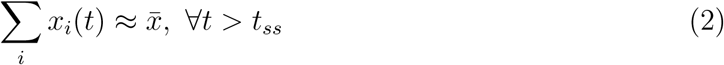

for some desired population size 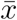. Differentiating equation (2) we get 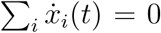 (note that this differs from the steady state of the system where 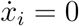 for all *i*) implying that

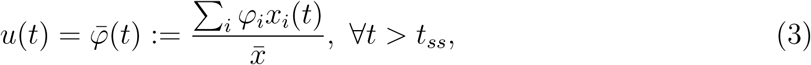

which is the weighted average growth rate.

Substituting (3) into (1) yields

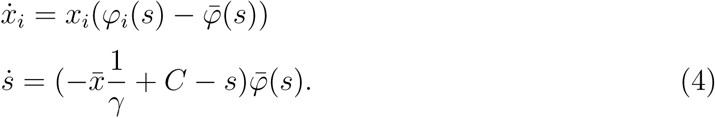

Furthermore, since cell population and substrate are a conserved quantity, we can restrict analysis to the attractive manifold 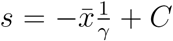, further simplifying our continuous model to

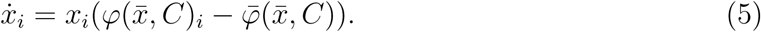

When considering basic bacterial growth in large (greater than 1000 individuals) populations of cells, mass action models such as (4) are accurate. However, when considering rare events such as mutation mass action models become inaccurate and stochastic models are preferable. To model growth dynamics and mutational dynamics together we use a stochastic hybrid systems(SHS) model (9). In this model, equation (4) comprises the continuous state of a SHS that has reset maps

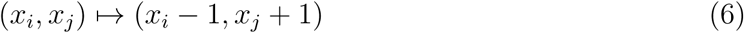

with transition intensities *λ*_*ij*_(*x*_*i*_). In other words, at rate *λ*_*ij*_(*x*_*i*_) a member of genotype *i* mutates to genotype *j*, removing one cell of type *i* and adding one cell of type *j*. In order to model *λ*_*ij*_(*x*_*i*_), we make the following assumptions:

1. Mutations are sufficiently rare that only individual mutants arise and either fix or are displaced (i.e., there is no clonal interference).
2. The population is large (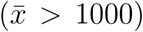). This is true for most microbial populations of interest, excluding some microfluidic devices.
3. The population size is constant.
4. Mutations happen at a constant rate per reproduction event.
5. Cell and nutrient mass are at steady state.

Assumptions 1 and 2 imply the standard strong selection weak mutation (SSWM) assumption used in literature (13). While assumptions 1 and 2 are stricter than the SSWM assumption, assumption 2 is motivated by the experimental setup we are considering and is therefore more reasonable then simply taking SSWM outright. Assumptions 2 and 3 imply that substitution happens via a Moran process and therefore the probability that a new mutant *i* fixes is a function of it’s fitness relative to the population average fitness 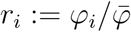,

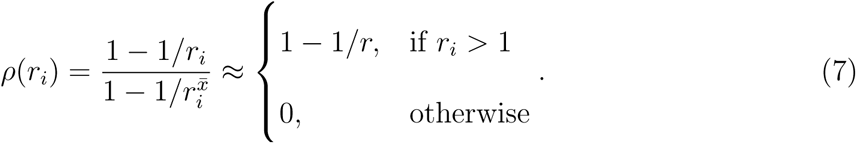

For more detail see Nowak Chapter 6 (14) and references within.

To compute the rate that new mutants appear, we suppose that the current population is homogeneous with fitness *φ*_1_. The chance a new mutant that will eventually fix in the population arises in the next Δ*t* seconds is proportional to the number of children born, *φ*_1_*x*_1_Δ*t*, times the chance it has of fixing, 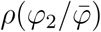. i.e.,

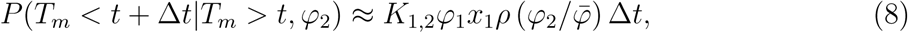

where *T*_*m*_ is the mutant’s time of arrival and *K*_1,2_ is some independent constant to represent proportionality as an equality. It follows from the definition of transition intensity(9) that

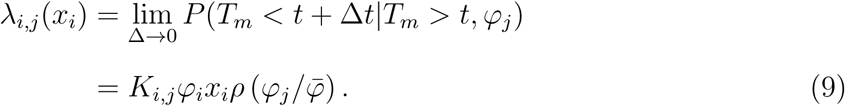

### Evolutionary Rate in a Turbidostat

With the continuous dynamics (4) and the transition intensities (9) we can now compute the time it takes a small sub-population with greater than average fitness to sweep the population.

Here, we define the *substitution time, T*_*s*_, as the time it takes a cell of genotype *i* to reach a population fraction of 50% starting from a single cell. This definition differs from what is typically used when considering a birth death processes. Since the dynamics are continuous, it is impossible to reach a population fraction of 1 in finite time when starting from an initial fraction of less than 1. Instead we choose a threshold where the novel cell genotype could be easily detected.

To compute the substitution time, we begin with our simplified continous model in equation (5), where s is treated as constant and cell densities are at steady state. We also note that *x* may be normalized by 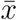, in which case units are population fraction. For the remainder of this paper we will use population fraction when referring to *x*.

Equation (5) has solution

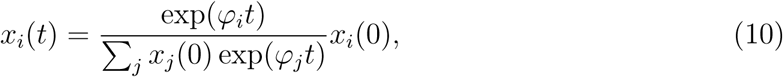

which for the case of genotype 2 superseding genotype 1, may be used to solve 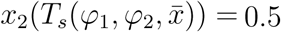 for 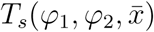) yielding

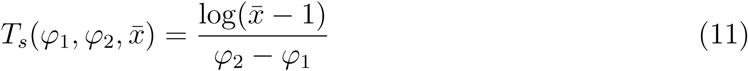

for 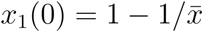, 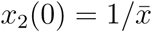, and *φ*_2_ > *φ*_1_.

Next, we compute the time to appearance of the next mutant *T*_*m*_, which is captured by a Poisson process(9). For a homogeneous population, as we have assumed, we can compute the expected time to mutation by

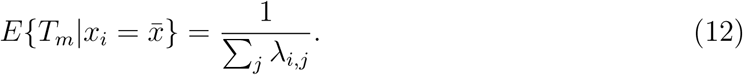

A turbidostat allows us to pick the population size 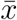, while a chemostat allows us to pick the average growth rate 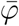. A turbidostat with mixture control enables us to control both 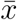 and 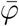 by choosing a fixed dilution rate and then adjusting the ratio of media sources until nutrient limitation is reached. This allows the population size 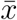 and the average growth rate 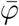 to be free variables. The time to appearance and substitution of a new mutant with fitness *φ*_2_ given a homogeneous population with fitness *φ*_1_ is

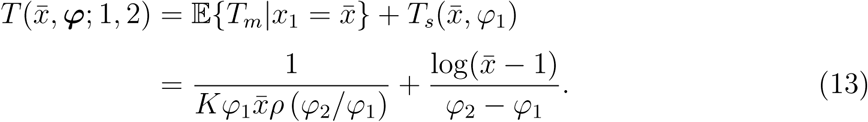

The time to appearance and substitution given in equation (13) has no closed form minimum over 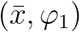 but may be optimized numerically (Fig. 1).

**Figure 1:**
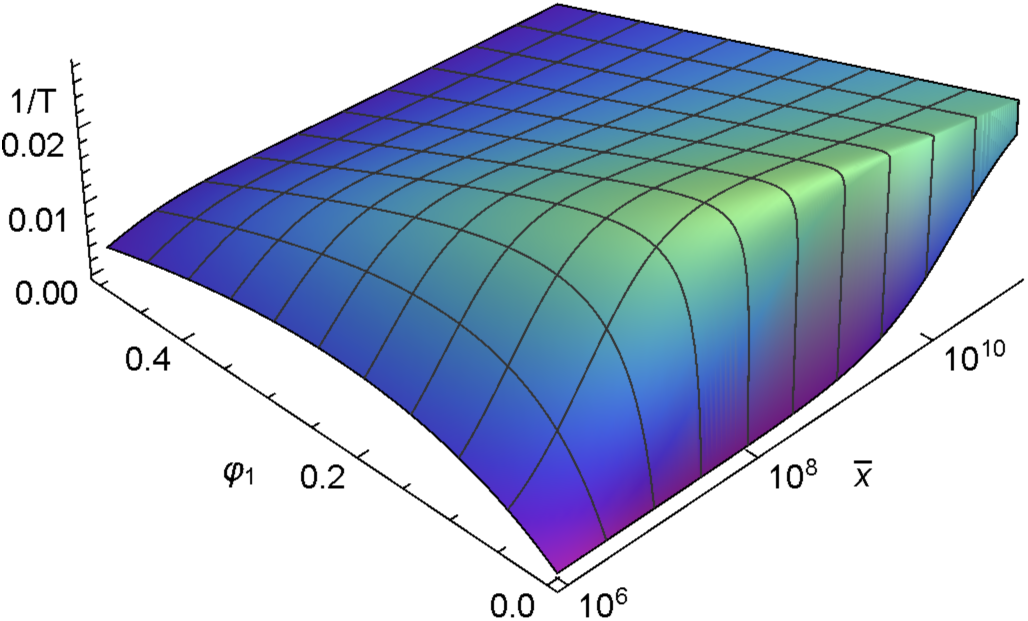
Rate of substitution (per hour) for a homogeneous population of genotype 1 to genotype 2 assuming k = 10^−7^ and *φ*_2_ = 0.6. For small 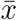 the penalty grows close to 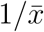 while for large 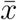 the penalty grows with 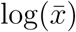.

### Circumventing Clonal Interference

We can now use our evolutionary model to explore a more complex landscape involving two alleles and negative epistasis (Fig 2). Applying our previous result to two substitution events with the optimal population size we get that the time to substitution of AB along the path ab → Ab → AB is 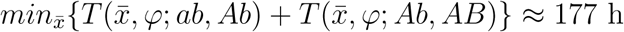 while the substitution time along the path ab → aB → AB is approximately 907 h. Clearly the former path is preferable from an engineering standpoint, however due to clonal interference, if phenotype aB arises at any point before phenotype AB, final fixation of AB will progress more slowly due to type aB’s superior growth rate with respect to type Ab and AB’s small improvement over aB (15). Indeed, regardless of weather AB arises from Ab or aB, the population dynamics (10) imply that the progress of type AB’s sweep relative to aB happens at a rate *φ*_*AB*_ − (*φ*_*aB*_*x*_*aB*_ − *φ*_*AB*_*x*_*AB*_)/(*x*_*aB*_ + *x*_*AB*_), which is very small for small differences growth rate and will delay the fixation of AB when the fraction *x*_*aB*_/*x*_*AB*_ is significant.

**Figure 2:**
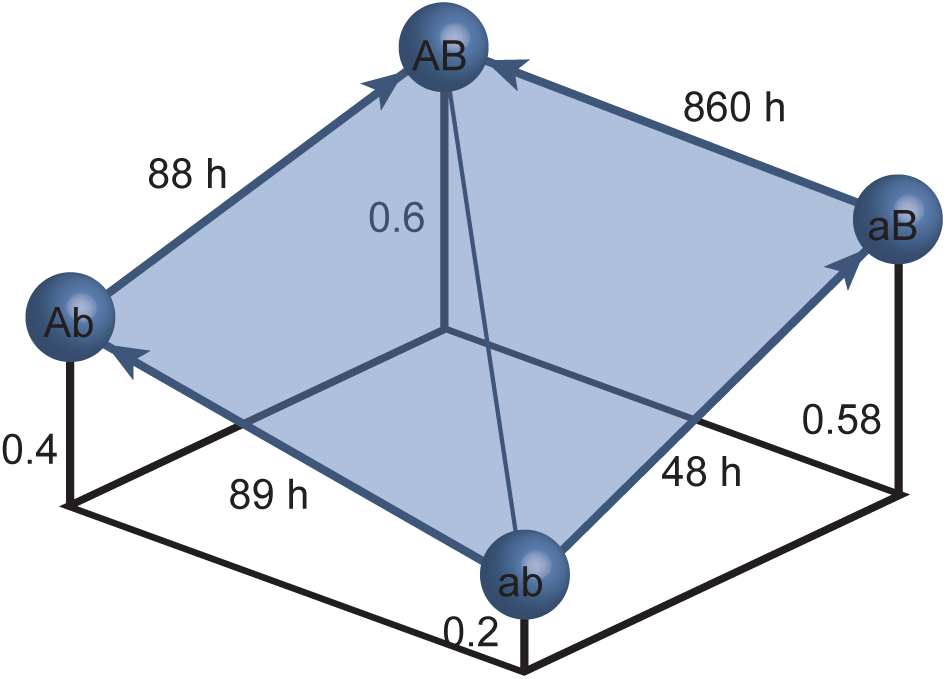
Example fitness landscape suffering from negative epistasis. In this example, two loci are considered each with two alleals, yielding four genotypes, ab, Ab, aB, and AB. The length of each “lolipop” denotes growth rate per hour. Edge weights and direction denote the expected substitution time for an optimal population size, given a mutation rate of 10^−7^ per cell division.

To explore conditions under which evolution is optimized in this environment we compute the probability that an evolutionary trajectory will progress along ab → Ab → AB, which can be approximated by

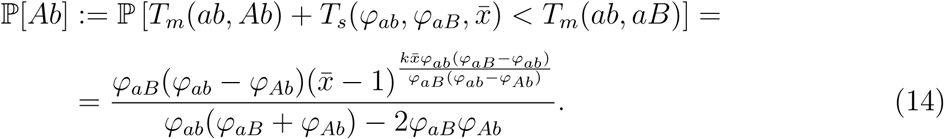

Equation (14) allows us to optimize the rate of two step evolution for known fitness landscapes. We get that the expected time to substitution of AB starting from ab is

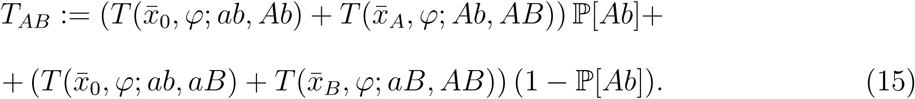

*T*_*AB*_ can be minimized over 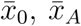 and 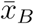 to find an optimal population size for each point in our hill climb. Applying this strategy to our example gives 837 h at an initial population size of approximately 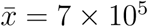.

### Rapid Evolution in an N-plex turbidostat

In the previous section, we showed that for the case of epistasis there is a fastest evolutionary path. For the case of negative episatis, clonal interference ensures that the probability that the fastest path will be taken is never more than 50%. Recent developments have made small multiplex bioreactors possible (8, 16) enabling parallel evolutions. With parallel bioreactors an experimenter only needs a single population of many to complete a hill climb before the evolved strain can be collected. This multiplex experimental setup then allows us to model the evolution of our meta-population (the population of individual communities within each bioreactor) as a N individual Bernoulli trials. For the two locus example, the probability of success (taking the faster path) is given in equation (14) and the probability that at least one of N cultures evolves along the fastest path is

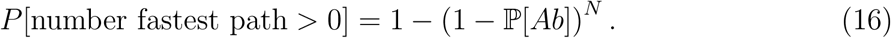

We can now subject equation (16) to either specification (e.g. Find 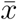 so that there is a 95% chance of at least one population taking the fastest path) or optimization. Fig. 3 shows the expected time and optimal per bioreactor population size as a function of number of bioreactors. Adding multiple bioreactors significantly decreases the wait time for an evolved strains with returns diminishing beyond 15 bioreactors.

**Figure 3:**
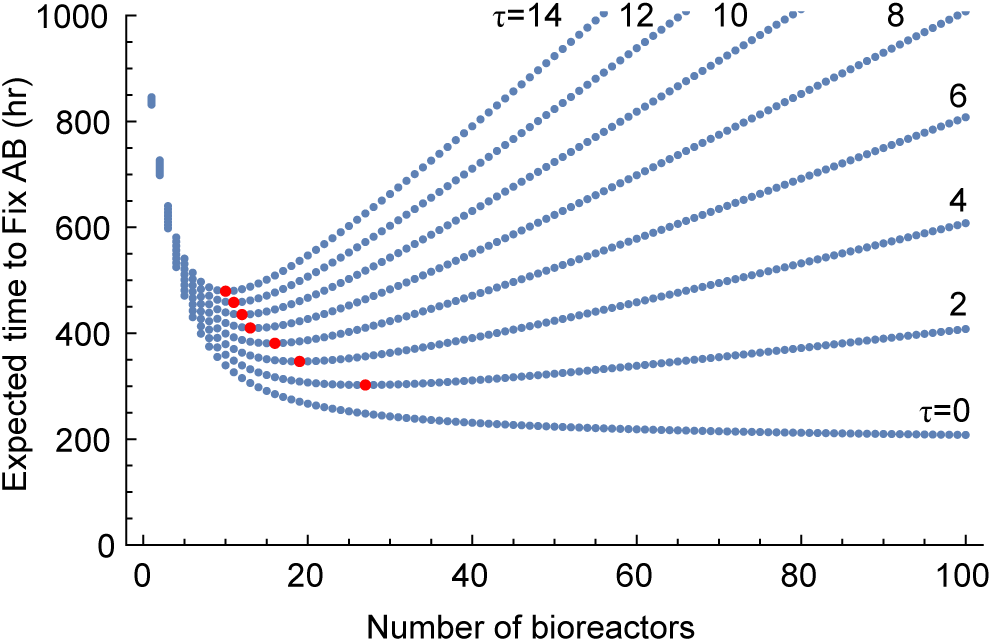
The relationship between the number of bioreactors and optimal rate of evolution (*τ* = 0). Plot corresponding to *τ* > 0 are given a linear penalty for the number of bioreactors, representing the additional labor required to setup and maintain additional bioreactors in units of culture time. For example, if each additional bioreactor takes 1 hour of labor and 1 hour of labor is worth 10 hours of waiting then *τ* = 1 × 10 hours per bioreactor. The points in red indicate the optimal number of bioreactors for each value of *τ*.

### A control strategy for parallel turbidostats

In prior sections, our optimization strategies have required exact knowledge of the fitness landscape. In practice this information is rarely known. Instead, estimates of optimal fitness may be available by applying knowledge of the metabolic load of expressing enzymes as well as the fitness benefit of downstream metabolites. With an estimate of the optimum fitness and insights gained from the previous sections, we can devise a heuristic control strategy to accelerate evolutionary hill climbs in parallel turbidostats.

We believe that the following heuristics apply in general and can be used to generate a control strategies that accelerate hill climbs of metapopulations.

1. Due to clonal interference, populations on the same order (or greater) as the mutation rate tend to take the largest fitness jumps available.
2. Larger jumps provide diminishing returns and should be avoided.

Using the stated heuristics, we have devised the control strategy outlined in Algorithm 1. To evaluate our control strategy we compared it’s performance in simulation to that of the null strategy where *N* turbidostats are inoculated but left without further manipulations.

#### Algorithm 1

1. Inoculate *N* turbidostats
2. *φ*_*m*_ ← growth rate of inoculum.
3. *φ*_*N*_ ← hypothesized maximal growth rate.
4. **while** *φ*_*m*_ < *φ*_*M*_ **do**
5. **for all** chambers *c* **do**
6. *φ*_*c*_ ← growth rate of chamber *c*
7. Δ ← *φ*_*M*_ − *φ*_*m*_
8. **if** *φ*_*M*_ − 0.25Δ > *φ*_*c*_ > *φ*_*m*_ + 0.25Δ **then**
9. Restart all chambers from chamber *c*
10. *φ*_*m*_ ← *φ*_*c*_

Our simulation implements the SHS with continuous state as in equation (5) and reset map given in equation (6) and terminates when a single chamber reaches 50% population fraction of the most fit type. We choose a randomly generated fitness landscape with three epistatically interacting loci (e.g. binding affinity, expression, and importation), each with a single alternate allele conferring a fitness benefit regardless of order of fixation (i.e. no sign episatis). Fig 4 shows the mean time reach the fitness peak for Algorithm 1 vs the null algorithm. Comparing Algorithm 1 with the null algorithm we see an improvement of nearly 40%. In addition to the improved optimal time, Algorithm 1 is also much more robust to the choice of an overly small population and is no worse than the null algorithm for large populations. The robustness of Algorithm 1 is an important feature because the a precise value for the optimal population size is not known a priori.

**Figure 4:**
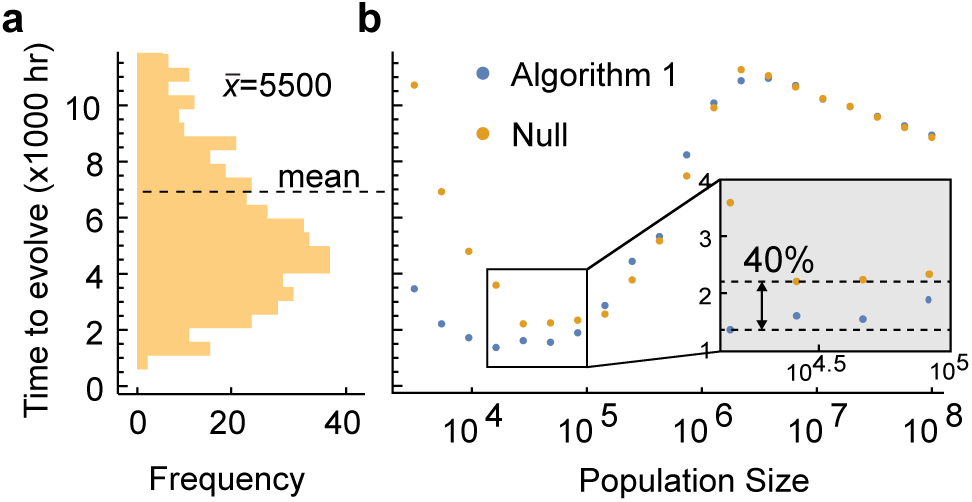
Performance of Algorithm 1 vs null algorithm for eight turbidostats. **a**, Simulated time to to reach fitness peak over 500 trials. **b**, Mean time to reach fitness peak of Algorithm 1 and the null algorithm for varying population sizes. Algorithm 1 still performs well when the population is chosen smaller than is optimal and performs no worse than null for large populations.

## Discussion

We introduce a simplified stochastic hybrid systems model that captures the general features of evolving populations in a turbidostat while remaining analytically tractable. With our model we show that the expected time to reach an evolutionary fitness peak can be computed analytically when the fitness landscape is known. From the analytical expression for expected time to hill climb we can optimize the population size to minimize the time to fixation of the most fit genotype. Our optimization then leads to a set of heuristics that can be used to devise control strategies to further optimize the time to evolve by directly manipulating meta-populations of turbidostats. We show that one such control method enables acceleration of evolution by nearly 40% while also making the evolutionary time more robust to choice of population.

Recent work has enabled the real world application of these control policies(8, 16, 17). Applications could include metabolic engineering problems where optimal growth rates can be extrapolated from known systems, and synthetic biology where it may be used to predict how “unhealthy” a cell may be made before it is likely to mutate in a given time span. Systems where engineered performance can be tied to metabolism may also benefit by showing experimenters how to optimize conditions, which in turn will increase research productivity.

In addition to direct applications, future work may focus on alternate algorithms for controlling meta-populations of turbidostats. Algorithms that allow a wider breath of evolutionary paths simultaneously may work better, especially in rugged landscapes. The application of stochastic remixing of populations may also provide more robust of faster control strategies. We believe that further exploration will reveal a rich space of control methods that have implications for both engineering and experimental evolution.

## References

1 Ofria, C., and Wilke, C. O. (2004) Avida: A Software Platform for Research in Computational Evolutionary Biology. Artificial Life 10, 191–229.

2 Ray, T. S. Synthetic life: Evolution and optimization of digital organisms. Scientific Excellence in Supercomputing: The 1990 IBM Contest Prize Papers. 1992; pp 489–531.

3 Kauffman, S., and Levin, S. (1987) Towards a general theory of adaptive walks on rugged landscapes. Journal of Theoretical Biology 128, 11–45.

4 Geritz, S., Kisdi, E., Meszena, G., and Metz, J. (1997) Evolutionary Ecology 12, 35–57.

5 Otto, S. P., and Day, T. Abiologist’s guide to mathematical modeling in ecology and evolution; Princeton University Press, 2007; Vol. 13.

6 Ro, D.-K., Paradise, E. M., Ouellet, M., Fisher, K. J., Newman, K. L., Ndungu, J. M., Ho, K. A., Eachus, R. A., Ham, T. S., Kirby, J., Chang, M. C. Y., Withers, S. T., Shiba, Y., Sarpong, R., and Keasling, J. D. (2006) Production of the antimalarial drug precursor artemisinic acid in engineered yeast. Nature 440, 940–943.

7 Raman, S., Rogers, J. K., Taylor, N. D., and Church, G. M. (2014) Evolution-guided optimization of biosynthetic pathways. Proceedings of the National Academy of Sciences 111, 17803–17808.

8 Takahashi, C. N., Miller, A. W., Ekness, F., Dunham, M. J., and Klavins, E. (2015) A Low Cost, Customizable Turbidostat for Use in Synthetic Circuit Characterization. ACS Synthetic Biology 4, 32–38.

9 Hespanha, J. P. (2006) Modelling and analysis of stochastic hybrid systems. IEE Pro-ceedings – Control Theory & Applications 153, 520–535.

10 Smith, H. L., and Waltman, P. The Theory of the Chemostat; Cambridge University Press, 1995.

11 Bryson, V., and Szybalski, W. (1952) Microbial selection. Science 116, 45–51.

12 Moran, P. A. P. The statistical processes of evolutionary theory; Clarendon Press, 1962.

13 Gillespie, J. H. (1983) Some Properties of Finite Populations Experiencing Strong Selection and Weak Mutation. The American Naturalist 121, pp. 691–708.

14 Nowak, M. A. Evolutionary Dynamics: Exploring the Equations of Life; Harvard University Press, 2006.

15 Gerrish, P., and Lenski, R. (1998) The fate of competing beneficial mutations in an asexual population. Genetica 102-103, 127–144.

16 Miller, A. W., Befort, C., Kerr, E. O., and Dunham, M. J. (2013) Design and Use of Multiplexed Chemostat Arrays. Journal of Visualized Experiments

17 Toprak, E., Veres, A., Michel, J. B., Chait, R., Hartl, D. L., and Kishony, R. (2012) Evolutionary paths to antibiotic resistance under dynamically sustained drug selection. Nat. Genet. 44, 101–105.

